# Cigarette smoke induces colon cancer by regulating the gut microbiota and related metabolites

**DOI:** 10.64898/2026.01.30.702732

**Authors:** Ya-Nan Bao, Quan-Ming Zhao, Xing-Hao Yang, Yi-Bo Gong, Bei-Bei Gan, Wen-Liang Li

## Abstract

The causal relationship between smoking and colorectal cancer (CRC) remains unclear. In this study, a cigarette smoke–exposed mouse model demonstrated that smoking significantly increased CRC incidence by inducing gut microbiota dysbiosis and altering related metabolites. Smoke exposure reduced beneficial bacteria (e.g., Lactobacillus), increased harmful bacteria (e.g., Firmicutes and Clostridium), elevated metabolites such as histamine, and suppressed the tumor suppressor genes PARG, CPT2, and ALDH1A1, thereby promoting tumor development. Functional assays in CRC cell lines further confirmed that CPT2 knockdown enhanced malignant phenotypes, including proliferation, migration, and invasion. Clinical analysis showed that these genes were markedly downregulated in smoking-related CRC patients, with strong diagnostic value (AUC > 0.8).

**Conclusion:** Smoking promotes CRC by inducing gut microbiota dysbiosis, metabolic reprogramming, and suppression of tumor suppressor genes, particularly CPT2. These findings highlight the importance of smoking cessation in CRC prevention and provide potential biomarkers for early diagnosis and therapeutic intervention.

## Introduction

Colorectal cancer (CRC) is one of the most common malignant tumors of the digestive tract. According to global cancer statistics, in 2020, the incidence rate of CRC ranked third, and its mortality rate ranked second [1]. Research has shown that CRC is a heterogeneous disease regulated by multiple factors and has a complex biological process. Its occurrence and development mechanisms still need to be studied. There is increasing evidence that lifestyle factors such as diet, smoking, obesity, and exercise are associated with CRC [3,4].

The occurrence and development of CRC is a heterogeneous process of changes in somatic cell molecules influenced by diet, the environment, microbial exposure, and host immunity. Most cases of CRC are not caused by familial genetic factors but rather are sporadic [5,6]. A person’s dietary habits can profoundly impact the initiation, promotion, and progression of tumors. Smoking increases the risk of lung cancer, with approximately 80% of primary lung cancer cases attributable to smoking [7]. Smoking also increases the risk of cancer in other organs that are not directly exposed to cigarette smoke, such as the colon, rectum, pancreas, and kidneys. Research shows that smoking is significantly related to the incidence and mortality rates associated with human CRC, and in animal models, smoking increases the risk of CRC development [4]. Smoking is currently a clear risk factor for CRC, which may be related to carcinogens such as polycyclic aromatic hydrocarbons and aromatic amines in tobacco smoke [8]. However, the mechanism by which smoking promotes the occurrence and progression of CRC remains unclear. An increase in bacterial diversity has been observed in humans after they quit smoking. A previous report also indicated that changes in the microbiome and mucin structure are associated with smoking [9]. Smoking directly leads to changes in the structure of the gut microbiota, which in turn affects the pathological and physiological processes of chronic diseases through these changes in gut microbiota metabolism. In addition, we have fully confirmed the association between changes in the gut microbiota and CRC. However, whether changes in the gut microbiota represent a link between smoking and CRC remains elusive.

This study aimed to investigate the factors related to the occurrence of CRC, smoking, and changes in the gut microbiota and to establish a reliable animal model to accurately assess the role of smoking in altering the gut microbiota in the occurrence and development of CRC. The role of smoking in the development of CRC was determined via conventional and sterile mouse models. Smoking can promote CRC by inducing dysbiosis in the gut microbiota, which affects metabolites and influences CRC occurrence and development. Translational medicine research for cancer prevention and treatment should be promoted to gain a deeper understanding of the molecular mechanisms underlying cancer occurrence and development in the later stages and obtain a solid foundation for the early, effective, and specific diagnosis and treatment of tumors.

## Materials and methods

### Construction of mouse colon cancer model ^[26]^

In this study, 46 SPF-grade male C57BL/6 mice, aged 10 weeks and weighing 20-25g, were purchased from Spbf (Beijing) Biotechnology Co., Ltd. The mice were divided into 3 groups: Group A (control group, n=6); Group B (GF-AOM group, n=20); and Group C (GF-AOMS group, n=20). Group B mice were intraperitoneally injected with the carcinogen AOM (10 mg/kg) once a week for 6 consecutive weeks to induce colorectal cancer (CRC). Group B mice were exposed to clean air for 2 hours daily, for 28 weeks; Group C mice were exposed to cigarette smoke (4% concentration) for 2 hours daily, for 28 weeks. The cigarette smoke was generated by a peristaltic pump, mixed with fresh air, and then pumped into the exposure chambers. After the experiment (week 28), after blood collection via the orbital venous plexus, the mice were anesthetized with 5% pentobarbital sodium (50 mg/kg). Following anesthesia, euthanasia was performed through cervical dislocation, and colon tissue samples were promptly harvested for subsequent analysis.

### HE staining

The colorectal tissues of each group of mice were fixed overnight with 4% paraformaldehyde (light recovery), dehydrated in ethanol (Chengdu Cologne Chemical Co., Ltd.) gradient, and then transparent in xylene (Xilong Chemical Co., Ltd.). After immersion in wax, they were embedded in paraffin and sliced at a thickness of 5 μ m. After staining with hematoxylin (Sevier) and eosin (Soleibao), the slides were sealed with neutral gum (Sevier) and randomly selected areas were observed under an optical microscope.

### IHC staining

Colorectal tissues from each group of mice were fixed overnight in 4% paraformaldehyde, dehydrated through graded ethanol, cleared in xylene, and embedded in paraffin after impregnation. The tissues were sectioned into 5 μm thick slices. After staining with hematoxylin and eosin, the sections were mounted with neutral resin and randomly selected regions were observed under an optical microscope.

### RT-PCR

Use TRIzol to isolate and purify RNA from the total sample according to the manufacturer’s provided operating protocol. Then, NanoDrop ND-1000 was used to control the quantity and purity of total RNA, and SureScript First strand cDNA synthesis kit was used at 25 ℃ for 5 minutes; 50℃, 15min; 85℃, 5s; At 4 ℃, the hold program reverse transcribes RNA into cDNA, dilutes cDNA 5-fold, and then adds detection factor primers(Table1) using a 5 × BlazeTaq qPCR Mix. Pre denature at 95 ℃ for 1 minute; Denaturation at 95 ℃ for 20 seconds; Annealing at 55 ℃ for 20 seconds; Extend the program at 72 ℃ for 30 seconds for 40 cycles of amplification, collect fluorescence and read CT values, and use the 2^- △△Ct^ method for data analysis.

### Western Blot

Use RIPA lysis buffer (Servicebio) containing protease inhibitor (Servicebio) at a concentration of 500ul to lyse colorectal tissue; Measure protein concentration using BCA protein assay kit (Biyun Tian); Then, the protein extracted from the tissue was separated using 10% SDS-PAGE and the SDS-PAGE gel was transferred onto a PVDF membrane (Millpore); Seal the membrane with 5% BSA (solarbio) and transfer the sealed PVDF membrane to the corresponding primary antibody for incubation; Subsequently, secondary antibody (Servicebio, 1:5000) incubation was carried out, and the highly sensitive ECL luminescence kit (Affinity) was used for color development. The Woote biochemical luminescence imaging analysis system was used for exposure, and images were collected. The grayscale values of the bands were obtained using the analysis software (Image J).

### Transcriptome sequencing、 Metabolome analysis and 16sRNA sequencing

To investigate the mechanism by which smoking exacerbates colorectal cancer, this study employed transcriptomic sequencing, metabolomic analysis, and 16S sequencing on an AOM-induced colorectal cancer mouse model. Mice from the Control group (A), GF-AOM group (B), and GF-AOMS group (C) were subjected to transcriptomic high-throughput sequencing, untargeted metabolomic sequencing, and 16S sequencing on their colorectal tissues, serum, and feces, respectively. Transcriptomic analysis included RNA extraction, quality control, sequence analysis, gene expression level assessment, and differential expression analysis to identify relevant genes. Metabolomic analysis involved calculating fold changes (FC), performing t-tests, and conducting PLS-DA analysis to identify differential metabolites and enrich metabolic pathways. 16S sequencing was used to compare the impact of smoking on gut microbiota and explore the role of high-abundance microbes in CRC.

### Association analysis

The 151 key genes from transcriptomics, 56 key metabolites from metabolomics, and 30 key microorganisms from the 16S microbiome were correlated. First, Spearman correlation was calculated between genes and metabolites, selecting those with |cor| > 0.6 and p < 0.05. Similarly, Spearman correlation between metabolites and microorganisms was calculated, selecting those with |cor| > 0.6 and p < 0.05. PPI network analysis was performed on genes related to metabolites, and the MCODE algorithm was used for further analysis to identify candidate genes. The expression of these candidate genes was examined using the GSE39582 colorectal cancer dataset from the GEO database. Finally, PCA, ROC, prognostic analysis, COX risk prediction, and combined diagnostic analysis were conducted on the key genes.

### Cell culture

Human normal colon epithelial cells (NCM460) and colorectal cancer cells (HT29) were purchased from Auragene (Changsha, China), while HCT116 and SW480 colorectal cancer cells were obtained from Procell (Wuhan, China). After recovery from liquid nitrogen, cells were maintained in RPMI-1640 complete medium (Meilunbio, MA0214) supplemented with 10% fetal bovine serum (FBS; Gibco, 10099-141) and 1% penicillin/streptomycin (Meilunbio, MA0110) at 37 °C in a humidified incubator with 5% CO₂. When cell confluence reached approximately 90%, cells were digested with 0.25% trypsin (Meilunbio, MA0233), washed with PBS, centrifuged at 1000 rpm for 5 min, and subcultured at a ratio of 1:2 into new flasks. For experiments, cells were seeded into 6-well plates at a density of 2.5–5 × 10⁵ cells/well, cultured for 24 h, and subsequently harvested for Western blot (WB) and quantitative real-time PCR (qPCR) analyses.

### Lentivirus packaging and infection

Lentiviral particles were produced in 293T cells by co-transfection with the transfer vector and packaging plasmids using PEI. Viral supernatants were collected at 48 and 72 h, clarified, and stored at −80 °C. HT29 cells were infected with the lentivirus in the presence of polybrene, followed by puromycin selection to establish stable cell lines. Knockdown efficiency of CPT2 was confirmed by Western blot and qPCR.

### Cell proliferation assay

Cell proliferation was assessed using the CCK-8 assay. Briefly, cells were seeded into 96-well plates (3–7 × 10³ cells/well) and cultured for 0, 24, 48, 72, and 96 h. At each time point, CCK-8 reagent was added and incubated for 0.5–4 h, and absorbance was measured at 450 nm.

### Cell invasion assay

Cell invasion was evaluated using a Transwell assay. Briefly, cells resuspended in serum-free medium were seeded into the upper chambers pre-coated with Matrigel, while medium containing serum was added to the lower chambers. After incubation at 37 °C for 12 h, invaded cells on the lower membrane surface were fixed, stained with crystal violet, and counted under a microscope.

### Cell invasion assay

Cell invasion was assessed using Transwell chambers coated with diluted ECM gel. Briefly, 2.5 × 10⁴ cells in serum-free medium were seeded into the upper chambers, and medium containing 10% FBS was added to the lower chambers. After incubation at 37 °C, invaded cells were fixed, stained with crystal violet, and counted in five random fields under a microscope.

### Apoptosis assay

Cell apoptosis was assessed by flow cytometry using Annexin V-PE/7-AAD staining. Briefly, collected cells were resuspended in binding buffer, stained with Annexin V-PE and 7-AAD, and incubated at room temperature in the dark. Samples were filtered to obtain single-cell suspensions, and at least 10,000 events per sample were analyzed by flow cytometry. Data were used to quantify viable, early apoptotic, late apoptotic, and necrotic cells.

### Cell cycle analysis

Cell cycle distribution was assessed by flow cytometry using propidium iodide (PI) staining. Briefly, collected cells were fixed in cold ethanol, treated with RNase A, and stained with PI. Samples were filtered to obtain single-cell suspensions, and at least 10,000 events per sample were analyzed by flow cytometry to determine the proportions of cells in G₀/G₁, S, and G₂/M phases.

### Statistical analysis

Statistical analysis was conducted using Prism Graphpad, and the experimental data were presented in the form of mean ± standard error (Mean ± SEM). T-test is used for inter group quantitative data comparison, and one-way analysis of variance is used for multi group quantitative data comparison. Setting p<0.05 has statistical significance: p<0.01 has significant statistical significance; P<0.001 has a very significant statistical difference; P<0.0001 has extremely significant statistical differences.

## Results

### AOM effectively induces colorectal tumorigenesis, and cigarette smoke further promotes tumor development

To investigate whether cigarette smoke promotes colorectal cancer (CRC), C57BL/6 mice were intraperitoneally injected with azoxymethane (AOM) for six consecutive weeks to induce CRC and subsequently exposed to either clean air or cigarette smoke. Body weight and fecal bleeding were monitored, and mice were sacrificed at 28 weeks(Fig.1A). Serum levels of LPS, TNF-α, and IL-6 were measured, and colons were collected to assess length, tumor incidence, epithelial thickness, and histopathology (H&E) and Ki67 expression (IHC).

**Figure 1.**
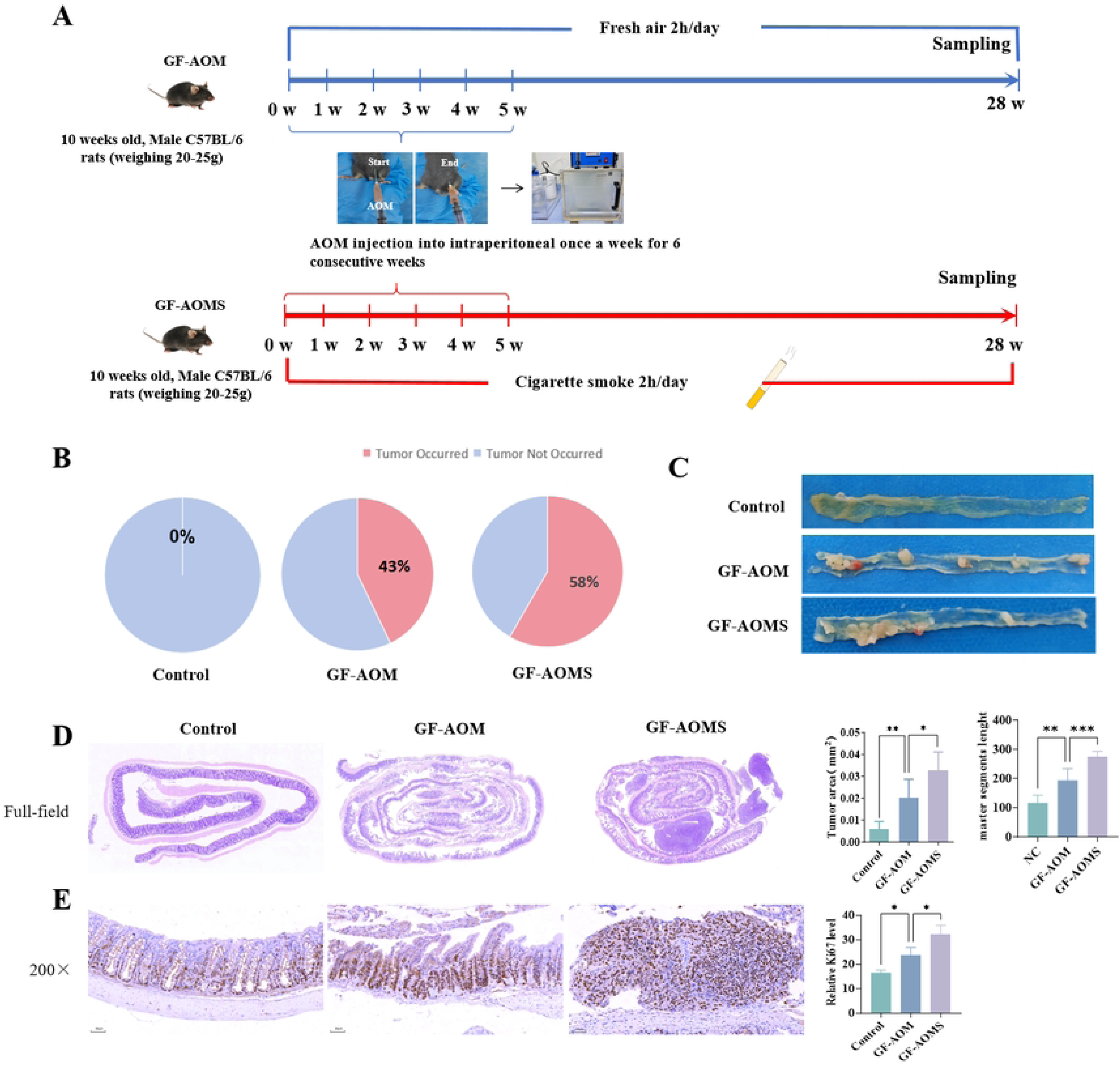
AOM and cigarette smoke promote colorectal tumorigenesis in mice. (A) Schematic of the experimental design. C57BL/6 mice were injected intraperitoneally with AOM to establish a CRC model and subsequently exposed to either fresh air or cigarette smoke for 2 h/day over 28 weeks. AOM (10 mg/kg) was administered once weekly for six consecutive weeks. Mice were sacrificed at week 28 (GF-AOM, n = 20; GF-AOMS, n = 20). (B) Tumor incidence in each group. (C) Representative images of colorectal tissues at the endpoint. (D) H&E staining of distal colorectal tissues showing epithelial hyperplasia and inflammatory infiltration; statistical analysis is shown (scale bar, 50 μm; magnification, 200×). (E) Immunohistochemical (IHC) staining of Ki67 in distal colorectal tissues; statistical analysis is shown (scale bar, 50 μm; magnification, 200×). Data are presented as mean ± SEM. Comparisons between two groups were performed using unpaired t-tests, and multiple group comparisons were performed using one-way ANOVA. *ns, p > 0.05; *p < 0.05; **p < 0.01; ***p < 0.001; ****p < 0.0001*.

Body weight analysis showed that Control mice gained weight steadily, whereas AOM-treated mice (GF-AOM) exhibited slowed weight gain compared with Control. Mice exposed to both AOM and cigarette smoke (GF-AOMS) displayed further weight reduction and greater fluctuations(sFig.1A). Fecal bleeding incidence increased from 0% in Control to 69% in GF-AOM and 92% in GF-AOMS mice(sFig.1B). Colon length was significantly shorter in GF-AOM mice than in Control (p < 0.0001), with a similar reduction observed in GF-AOMS mice(sFig.1C). Serum L-6(sFig.2A), TNF-α(sFig.2B) and LPS(sFig.2C), Ilevels were elevated in GF-AOM mice (p < 0.01) and further increased in GF-AOMS mice (p < 0.01), indicating compromised gut barrier function and enhanced systemic inflammation in the tumor microenvironment These results confirm that AOM successfully induces CRC in mice and that cigarette smoke exacerbates pathological phenotypes such as weight loss and fecal bleeding.

Gross examination(Fig.1C), H&E staining(Fig.1D), and IHC analysi revealed increased tumor incidence in AOM-treated mice compared with Control. Tumor formation was absent in Control mice, whereas GF-AOM mice exhibited a 43% tumor incidence with significantly increased tumor area (p < 0.01) and epithelial thickness (p < 0.01)(Fig.1B). Ki67-positive cells were also significantly elevated (p < 0.05)s(Fig.1E), indicating enhanced cell proliferation. In GF-AOMS mice, tumor incidence further increased to 58%, accompanied by larger tumor areas (p < 0.05), further epithelial thickening (p < 0.001), and higher Ki67-positive cell counts (p < 0.05). These findings indicate that cigarette smoke further promotes colorectal tumor growth in AOM-induced CRC mice.

### Multi-omics integration identifies seven smoking–CRC-associated key regulatory genes

To systematically elucidate the molecular mechanisms by which cigarette smoke synergizes with AOM to promote colorectal cancer (CRC), transcriptomic, fecal microbiome, and serum metabolomic analyses were performed on control (Control, A), AOM-treated (GF-AOM, B), and AOM plus cigarette smoke-exposed mice (GF-AOMS, C). Multi-omics integration allowed a comprehensive comparison of molecular alterations among groups.

### Transcriptomics

Differential gene expression analysis revealed 1,293 genes between B and A, enriched in 48 significant pathways (p < 0.05); 65 genes differed between C and B, enriched in 2 pathways (p < 0.05); and 1,085 genes differed between C and A, enriched in 52 significant pathways, including ribosome, PPAR signaling, ferroptosis, and NF-κB signaling (Fig. 3).

**Figure 2.**
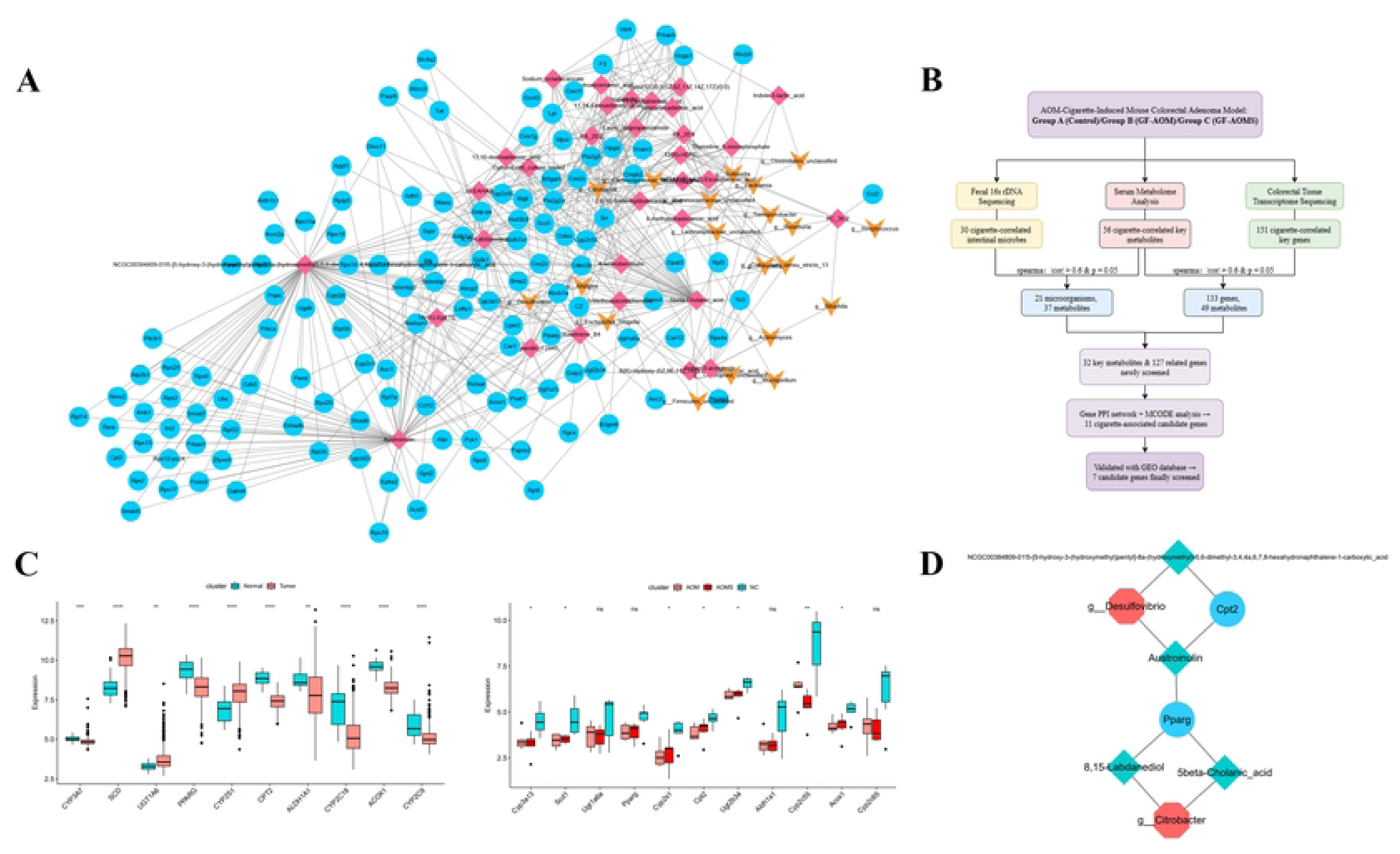
Smoking-associated gut microbiota–metabolite–key gene interactions and their diagnostic and prognostic relevance in colorectal cancer. (B) Key genes identified through integrated multi-omics analysis. (C) Boxplots of candidate gene expression in the GSE39582 dataset (left) and transcriptomic data (right). (D) PPI network illustrating interactions among key genes, metabolites, and microbes. (E) Schematic workflow of the integrated multi-omics analysis.

**Figure 3.**
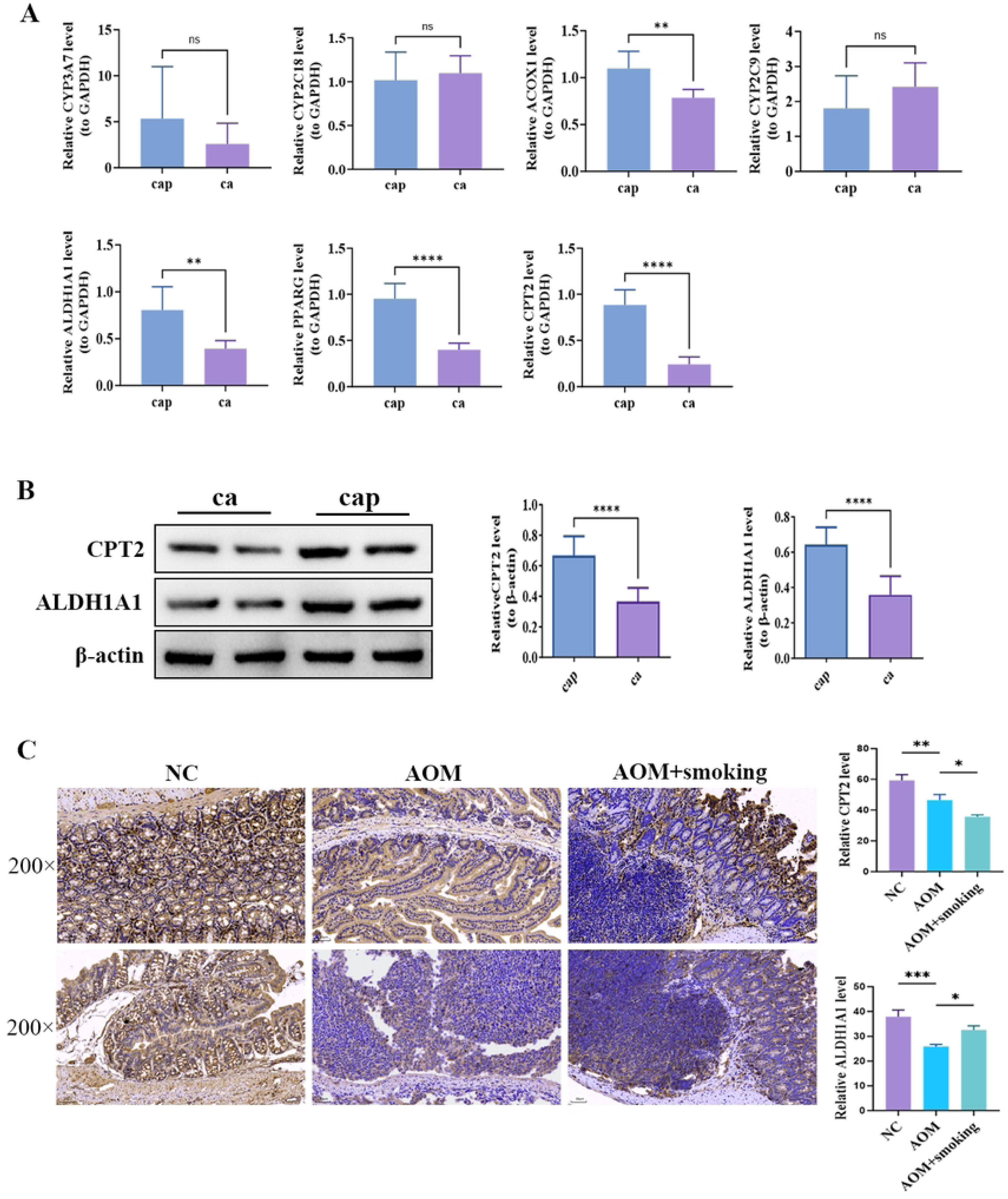
Validation of key gene expression in clinical and mouse samples. (A) qPCR analysis of seven candidate genes. (B) Western blot analysis of CPT2 and ALDH1A1 protein levels, using β-actin as a loading control. (C) IHC staining of CPT2 and ALDH1A1 in mouse colorectal tissues. Images were captured at 200× magnification; scale bar = 50 μm. Nuclei are stained blue, and CPT2/ALDH1A1-positive signals appear brown. (D) Clinical correlation analysis. Tissue types: ca, tumor; cap, adjacent normal tissue. Mouse groups: NC, control; AOM, AOM-treated; AOM+smoking, AOM plus cigarette smoke exposure. ImageJ Pro Plus was used for quantification, and GraphPad Prism was used for plotting. Data are presented as mean ± SEM. Statistical comparisons between two groups were performed using unpaired t-tests, and multiple groups were analyzed by one-way ANOVA. Significance levels: *ns, p > 0.05; *p < 0.05; **p < 0.01; ***p < 0.001; ****p < 0.0001*.

### Metabolomics

Non-targeted metabolomics identified 89 differential metabolites between B and A (34 enriched pathways), 24 metabolites between C and B (2 pathways), and 135 metabolites between C and A (73 pathways), implicating arachidonic acid metabolism, protein digestion and absorption, PPAR signaling, and TRP channel regulation by inflammatory mediators (Fig. 4).

**Figure 4.**
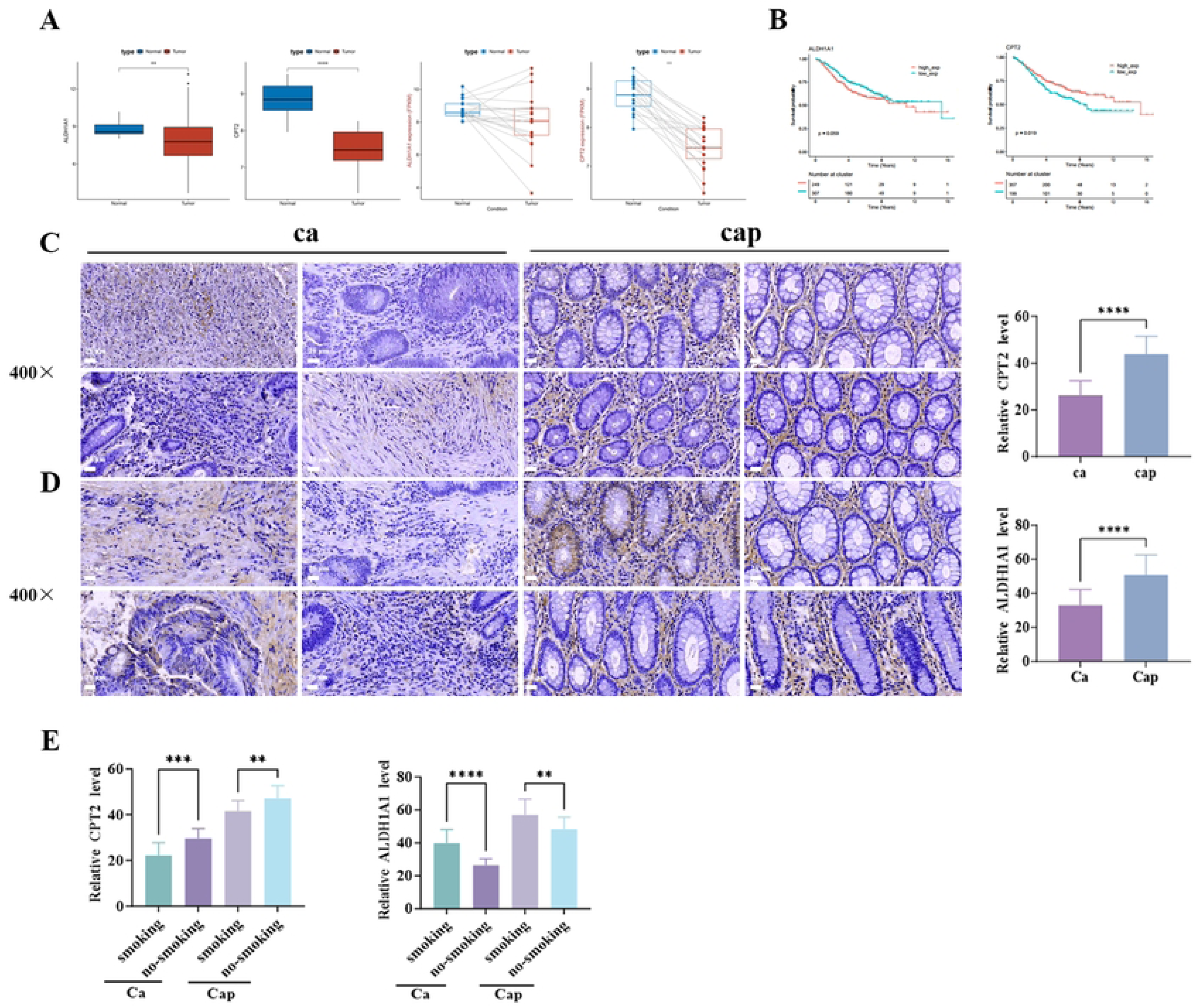
Expression patterns, survival association, and smoking-related effects of CPT2 and ALDH1A1 in colorectal cancer. (A) Boxplots showing differential expression of CPT2 and ALDH1A1 in tumor (ca) and adjacent normal tissues (cap). (B) Kaplan–Meier survival curves for CPT2 and ALDH1A1 in CRC patients. (C–D) IHC detection of CPT2 (C) and ALDH1A1 (D) in clinical CRC specimens; magnification 400×, scale bar = 25 μm. Nuclei are stained blue, and CPT2/ALDH1A1-positive signals are brown. (E–F) Impact of patient smoking status on CPT2 and ALDH1A1 expression based on IHC analysis. Data are presented as mean ± SEM. Comparisons between two groups were performed using unpaired t-test, and multiple group comparisons were assessed by one-way ANOVA. Statistical significance is indicated as *ns, p > 0.05; *p < 0.05; **p < 0.01; ***p < 0.001; ****p < 0.0001*.

### Microbiome

16S rDNA sequencing and alpha/beta diversity analyses indicated higher alpha diversity in the C group, with minimal clustering differences among groups. Firmicutes and Bacteroidetes were dominant phyla. Compared with A, B showed increased Bacteroidetes and Akkermansia; compared with B, C had higher Firmicutes and increased genera including NK4A136_group. LEfSe identified multiple differential taxa, ultimately yielding 30 smoking-related key microbial genera. Notably, smoking reduced beneficial bacteria such as Lactobacillus, while increasing Firmicutes, Mucispirillum, Clostridium, and Clostridioides, potentially disrupting the gut barrier and promoting tumorigenesis (Fig. 5).

**Figure 5.**
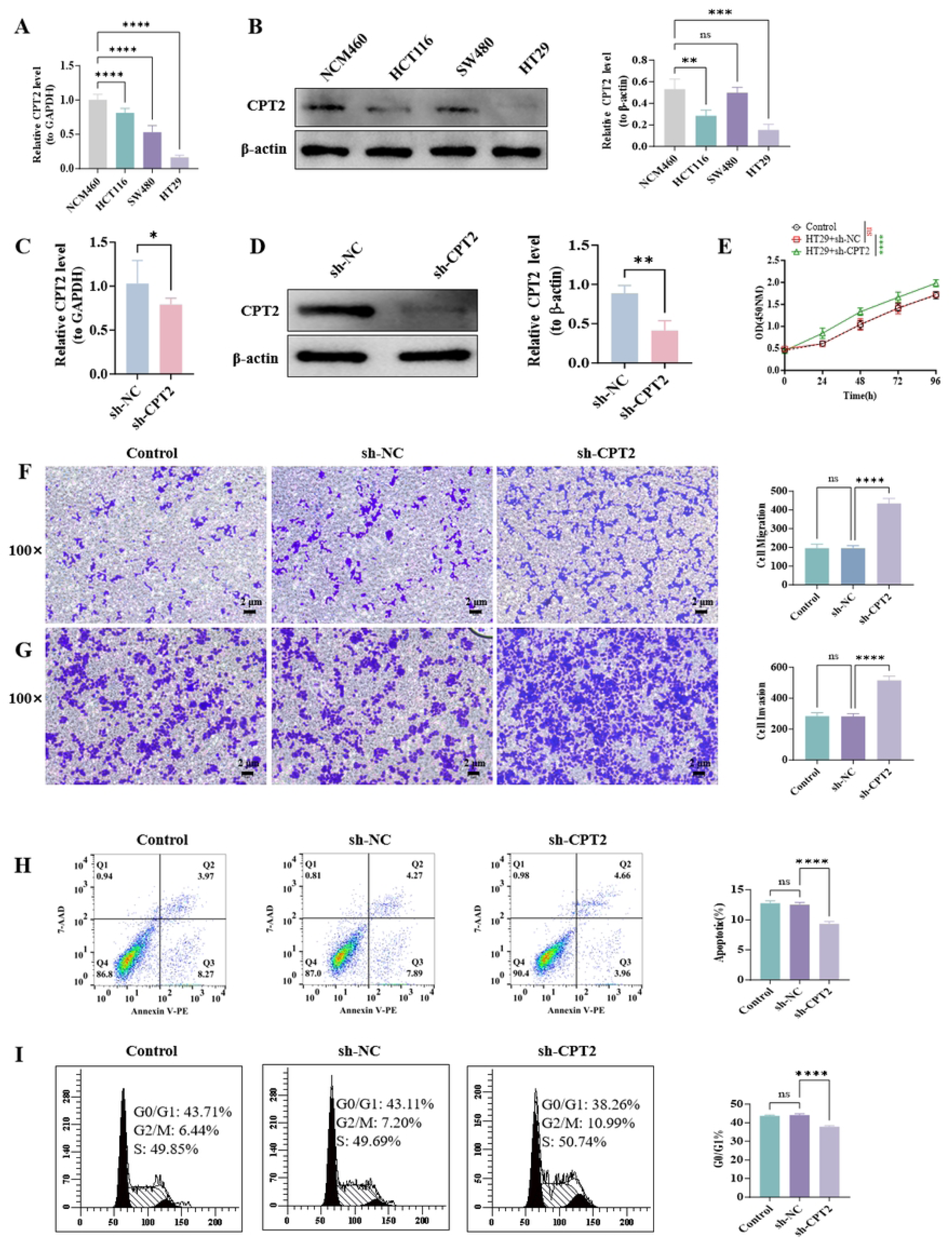
CPT2 knockdown promotes malignant phenotypes in HT29 cells. A: qPCR analysis of CPT2 mRNA expression in colorectal cancer cell lines (SW480, HCT116, HT29) and normal human colon mucosal epithelial cells (NCM460). B: Western blot analysis of CPT2 protein expression in SW480, HCT116, HT29, and NCM460 cells, with β-actin as the loading control. C: qPCR validation of CPT2 knockdown efficiency in HT29 cells. D: Western blot validation of CPT2 knockdown efficiency in HT29 cells, with β-actin as the loading control. E: CCK8 assay for cell proliferation. F: Transwell migration assay to assess cell migratory capacity. G: Matrigel invasion assay to evaluate cell invasive capacity. H: Flow cytometry analysis of cell apoptosis. I: Flow cytometry analysis of cell cycle distribution. Control represents HT29 cells without transduction, sh-NC indicates cells transduced with scrambled control vector, and sh-CPT2 represents HT29 cells with stable CPT2 knockdown. Data are presented as mean ± SEM. Comparisons between two groups were performed using unpaired t-tests, and multiple group comparisons were analyzed by one-way ANOVA. Statistical significance: *ns, p > 0.05; * p < 0.05; ** p < 0.01; *** p < 0.001; **** p < 0.0001*.

### Integration of multi-omics data

Venn and KEGG analyses of transcriptomic and metabolomic data between C and A groups identified 151 key genes, 56 key metabolites, and 30 key microbes. Association analyses revealed 667 gene–metabolite pairs (133 genes, 49 metabolites), 113 metabolite–microbe pairs (21 microbes, 37 metabolites), and 32 shared metabolites linked to 127 genes. PPI network analysis of these 127 genes identified 11 candidate genes (Cyp3a13, Scd1, Ugt1a6a, Pparg, Cyp2s1, Cpt2, Ugt2b34, Aldh1a1, Cyp2c55, Acox1, Cyp2c65). Validation in human CRC dataset GSE39582 and sequencing data refined this list to seven key genes: CYP3A7, PPARG, CPT2, ALDH1A1, CYP2C18, ACOX1, and CYP2C9 (Fig. 2B).

### Clinical relevance

PCA confirmed distinct separation of normal and CRC samples based on the seven-gene signature. ROC analysis showed high predictive performance for all candidate genes (AUC > 0.7). Diagnostic nomograms combining gene expression demonstrated high accuracy and net benefit, supported by decision curve and clinical impact analyses. Prognostic Cox models predicted 1-, 3-, and 5-year survival with high accuracy, and KM survival analysis indicated significant associations of some candidate genes with patient outcomes. Overall, PPARG and CPT2 showed strong diagnostic potential and relevance in CRC progression.

### Mechanistic associations

PPI network analysis of two highly correlated genes (PPARG, CPT2), four key metabolites (Austroinulin;NCGC00384809-01!5-[5-hydroxy-3-(hydroxymethyl) (A) pentyl]-8a-(hydroxymethyl)-5,6-dimethyl-3,4,4a,6,7,8-hexahydronaphthalene-1-carboxylic acid; 5 beta-Cholanic acid; 8,15-Labdanediol), and two pathogenic gut microbes (g_Desulfovibrio and g_Citrobacter) revealed direct interactions, suggesting that smoking may induce CRC through coordinated regulation of these factors (Fig. 2C).

### CPT2 and ALDH1A1 are downregulated in colorectal cancer and modulated by cigarette smoke

Based on integrated transcriptomic, metabolomic, and microbiome analyses of control (Control), AOM-induced CRC model (GF-AOM), and AOM plus cigarette smoke-exposed mice (GF-AOMS), seven candidate genes closely associated with CRC development were identified. To validate their clinical relevance, colorectal cancer and adjacent normal tissue samples were collected from patients for qPCR and Western blot (WB) analyses. To assess the effect of cigarette smoke on these key genes, mouse colorectal tissues were subjected to immunohistochemistry (IHC) staining.

qPCR analysis (Fig. 3A) revealed that ACOX1, CPT2, PPARG, and ALDH1A1 were significantly downregulated in CRC tissues compared with adjacent normal tissues (p < 0.01), whereas CYP3A7, CYP2C18, and CYP2C9 showed no significant differences. Literature indicates that ACOX1 regulates CRC proliferation and migration, while CPT2 and ALDH1A1 may modulate oxidative stress to affect tumor biology, though their mechanisms remain unclear. WB validation (Fig. 3B) confirmed that protein levels of CPT2 and ALDH1A1 were markedly lower in CRC tissues than in adjacent normal tissues (p < 0.0001).

IHC results (Fig. 3C) showed that, compared with Control mice, GF-AOM mice exhibited significantly reduced CPT2 and ALDH1A1 expression in colorectal tissues (p < 0.01). Notably, cigarette smoke exposure further decreased CPT2 expression in GF-AOMS mice (p < 0.05), whereas ALDH1A1 expression was slightly increased in the same group (p < 0.05). Clinical correlation analysis of CPT2 (Fig. 3D) indicated that its expression was significantly associated with sex (P = 0.003), with a higher proportion of females in the high-expression group, and smoking status (P = 0.008), with 100% of high-expression patients being non-smokers. No significant associations were observed between CPT2 expression and age, alcohol consumption, or TNM stage (P > 0.05).

These findings suggest that CPT2 and ALDH1A1 may serve as potential diagnostic markers or therapeutic targets in CRC, and that cigarette smoke exposure may modulate their expression in colorectal tumors.

### CPT2 and ALDH1A1 are downregulated in colorectal cancer and associated with patient prognosis

To further investigate the regulatory effect of smoking on CPT2 and ALDH1A1 in colorectal cancer (CRC), we analyzed their expression patterns and prognostic relevance in tumor and adjacent normal tissues from smoking and non-smoking patients.

Analysis of the human CRC dataset GSE39582 revealed that CPT2 and ALDH1A1 expression levels were significantly lower in tumor tissues compared to adjacent normal tissues (Fig. 4A), suggesting that downregulation of these genes may contribute to CRC development.

Kaplan–Meier survival analysis indicated that low CPT2 expression was associated with significantly reduced overall survival (OS) in CRC patients (p < 0.05), whereas high CPT2 expression correlated with better prognosis. In contrast, ALDH1A1 expression showed no statistically significant association with patient survival (p = 0.059) (Fig. 4B).

IHC analysis of clinical CRC specimens confirmed these findings, showing markedly reduced CPT2 (p < 0.001) and ALDH1A1 (p < 0.0001) expression in tumor tissues relative to adjacent normal tissues (Fig. 4C–D). Stratification by patient smoking status revealed that smoking further decreased CPT2 expression, whereas ALDH1A1 expression exhibited an opposite trend (Fig. 4E).

Overall, CPT2 and ALDH1A1 are both downregulated in CRC, with CPT2 low expression significantly associated with poor patient prognosis. These results suggest that smoking may exacerbate CRC progression by further suppressing CPT2 expression, highlighting CPT2 as a potential prognostic biomarker for CRC patients.

### CPT2 knockdown promotes malignant phenotypes in colorectal cancer cells with low endogenous CPT2 expression

To investigate the biological function of CPT2 in colorectal cancer (CRC), we screened CRC cell lines for low endogenous CPT2 expression and established stable CPT2 knockdown cell lines to assess its impact on tumor cell behavior. Initially, qPCR and Western blot were performed to determine CPT2 expression levels across three human CRC cell lines (SW480, HCT116, HT29) and a normal human colon mucosal epithelial cell line (NCM460). qPCR results showed significantly lower CPT2 mRNA levels in SW480, HCT116, and HT29 cells compared with NCM460 cells (p < 0.0001, Fig. 5A). Western blot analysis revealed significantly reduced CPT2 protein expression in HCT116 (p < 0.01) and HT29 cells (p < 0.001), whereas SW480 showed no significant difference (Fig. 5B). Based on these findings, HT29 cells were selected for subsequent experiments.

Stable CPT2 knockdown in HT29 cells was achieved via lentiviral transduction. qPCR and Western blot validation confirmed successful knockdown, with sh-CPT2 cells showing significantly decreased CPT2 mRNA (p < 0.05, Fig. 5C) and protein levels (p < 0.01, Fig. 5D) compared with sh-NC controls.

Functional assays were then conducted to evaluate malignant phenotypes. CCK8 assays demonstrated significantly enhanced proliferation in HT29+sh-CPT2 cells (p < 0.0001, Fig. 5E). Transwell migration (Fig. 5F) and Matrigel invasion assays (Fig. 5G) showed markedly increased migratory and invasive capacities in CPT2-knockdown cells (p < 0.0001). Flow cytometry analysis indicated reduced apoptosis (p < 0.0001, Fig. 5H) and alterations in cell cycle distribution, with decreased G0/G1 phase and increased S and G2/M phase proportions (Fig. 5I). In contrast, HT29+sh-NC cells showed no significant differences compared with control cells in proliferation, migration, invasion, apoptosis, or cell cycle distribution.

Collectively, these data indicate that CPT2 acts as a tumor suppressor in CRC. Its downregulation in HT29 cells promotes malignant phenotypes by enhancing proliferation, migration, and invasion, inhibiting apoptosis, and accelerating cell cycle progression. These findings suggest that decreased CPT2 expression may contribute to CRC progression by facilitating tumor cell growth and aggressive behavior.

## Discussion

In recent years, with the advancement of sequencing technology, researchers have increasingly focused on the effects of diet, smoking, alcohol consumption, and other lifestyle factors on the human gut microbiota. Studies have shown that the microbial community composition can rapidly change with alterations in diet, smoking, and alcohol intake [10]. Rodent microbiota transplantation experiments revealed that these changes can lead to gut microbiota dysbiosis, resulting in inflammation and metabolic disorders [11]. However, alterations in the gut microbiota do not necessarily constitute a prerequisite for functional changes. Smoking can modulate microbial metabolism, which in turn significantly impacts the host. The gut microbiota produces metabolites that interact synergistically or antagonistically, affecting key signaling pathways, including those related to metabolism and inflammation [12]. For instance, both human and animal studies indicate that the pathogenicity of microbes can be enhanced under changing environmental conditions, producing proinflammatory effects [13,14].

In this study, an AOM-induced colorectal tumor model in mice, combined with cigarette smoke exposure, was established. Pathological staining and Ki67 immunohistochemistry confirmed that smoking is a risk factor for CRC. While previous studies have shown an association between gut microbiota and smoking, it remains unclear whether microbiota changes contribute directly to CRC progression. To address this, we performed experiments combining AOM-induced CRC and cigarette smoke interventions to alter the gut microbiota in mice.

Our results revealed significant changes in microbial phyla, including a decreased Firmicutes-to-Bacteroidetes ratio and increased abundance of Firmicutes, Spirogyra, and Clostridium following long-term cigarette smoke exposure. Pathogenic species such as Clostridium difficile, C. perfringens, and C. botulinum can produce toxins that impair the intestinal barrier and promote tumorigenesis. These findings suggest that the Firmicutes/Clostridium ratio could serve as a potential indicator for gut microbiota health and CRC risk assessment.

Metabolomics analysis showed that cigarette smoke altered 13 intestinal metabolites, including histamine, 4-hydroxycinnamic acid, and 4-aminobenzoate, which are involved in 47 signaling pathways, such as arginine metabolism, tryptophan metabolism, arachidonic acid metabolism, and protein synthesis. Histamine, a chronic inflammatory mediator, can enhance inflammatory responses, promote immune escape, and remodel the tumor microenvironment [15]. Amino acid metabolism pathways, including arginine and tryptophan metabolism, play critical roles in immune modulation under stress, trauma, or tumor conditions [16–18].

Integration of transcriptomic, metabolomic, and microbiome data revealed that PPARG and CPT2, together with four metabolites (Austroinulin, NCGC00384809-015, [5-hydroxy-3-(hydroxymethyl)pentyl]-8a-(hydroxymethyl)-5,6-dimethyl-3,4,4a,6,7,8, 8-hexahydronaphthoxene-1-carboxylic acid, 5-beta-cholanic acid, and 8,15-labanediol) and two harmful gut bacteria (Desulfovibrio and Citrobacter), were significantly enriched in pathways associated with CRC progression. Both PPARG and CPT2 are highly expressed and can serve as diagnostic markers for CRC. PPARG, a nuclear receptor, regulates adipogenesis, lipid metabolism, insulin sensitivity, vascular homeostasis, and inflammation [19,20], and its dysregulation has been linked to carcinogenesis [21]. CPT2, a key enzyme in fatty acid oxidation, catalyzes the conversion of acylcarnitine to acyl-CoA [22,23]. Reduced CPT2 expression leads to acylcarnitine accumulation, which is associated with tumorigenesis [24].

In addition, our study extended these findings to cellular models. We selected HT29 cells, which have low endogenous CPT2 expression, to generate a stable CPT2 knockdown cell line. Functional assays demonstrated that CPT2 knockdown promoted malignant phenotypes, including accelerated proliferation, enhanced migration and invasion, decreased apoptosis, and altered cell cycle progression (reduced G0/G1 phase and increased S and G2/M phases). These effects are consistent with the pathway analyses showing that CPT2 regulates fatty acid oxidation, energy metabolism, and ROS homeostasis, which collectively influence CRC cell survival, proliferation, and invasiveness. Thus, CPT2 downregulation may contribute directly to CRC progression by promoting metabolic reprogramming and malignant cellular behaviors.

Clinically, PPARG and CPT2 were downregulated in tumor tissues compared to adjacent normal tissues. Cigarette smoke further suppressed their expression in both tumor and adjacent tissues, consistent with mouse sequencing and immunohistochemical results. Bioinformatic analyses indicated that low PPARG and CPT2 expression correlated with poor prognosis in CRC patients, suggesting their potential as protective prognostic biomarkers.

In conclusion, cigarette smoke promotes CRC development by inducing gut microbiota dysbiosis, leading to decreased PPARG and CPT2 expression and altered metabolite profiles. CPT2 knockdown experiments further confirmed that reduced CPT2 promotes malignant phenotypes in CRC cells via metabolic and signaling pathways. Together, PPARG and CPT2 represent candidate prognostic biomarkers and potential therapeutic targets for smoking-associated CRC.

## RESOURCE AVAILABILITY

### Lead contact

Further information and requests for resources and reagents should be directed to and will be fulfilled by the lead contact, [Ya-Nan Bao] ().

### Materials availability

This study did not generate new unique reagents.

## AUTHOR CONTRIBUTIONS

Y.-N. Bao and Q.-M. Zhao contributed to the study conceptualization and design. W.-L. Li supervised the study and served as Lead Contact. X.-H. Yang and Y.-B. Gong contributed to the methodology. Data curation, investigation, validation, visualization, writing – original draft, and writing – review & editing were performed mainly by B.-B. Gan. X.-H. Yang, Y.-B. Gong, and Y.-N. Bao helped carry out the experiments. Writing – review & editing were performed by Y.-N. Bao and Q.-M. Zhao. All authors read and approved the final manuscript. Y.-N. Bao and Q.-M. Zhao contributed equally. Y.-N. Bao and Q.-M. Zhao are Senior Authors. W.-L. Li served as Lead Contact.

## ACKNOWLEDGEMENTS

This work was supported by the Scientific Research Project of Yunnan Clinical Medicine Center (2024YNLCYXZX0415) and the Key Research and Development Program (202403AC100001).

## Supplementary Figure Legends

**Supplementary Figure 1.**
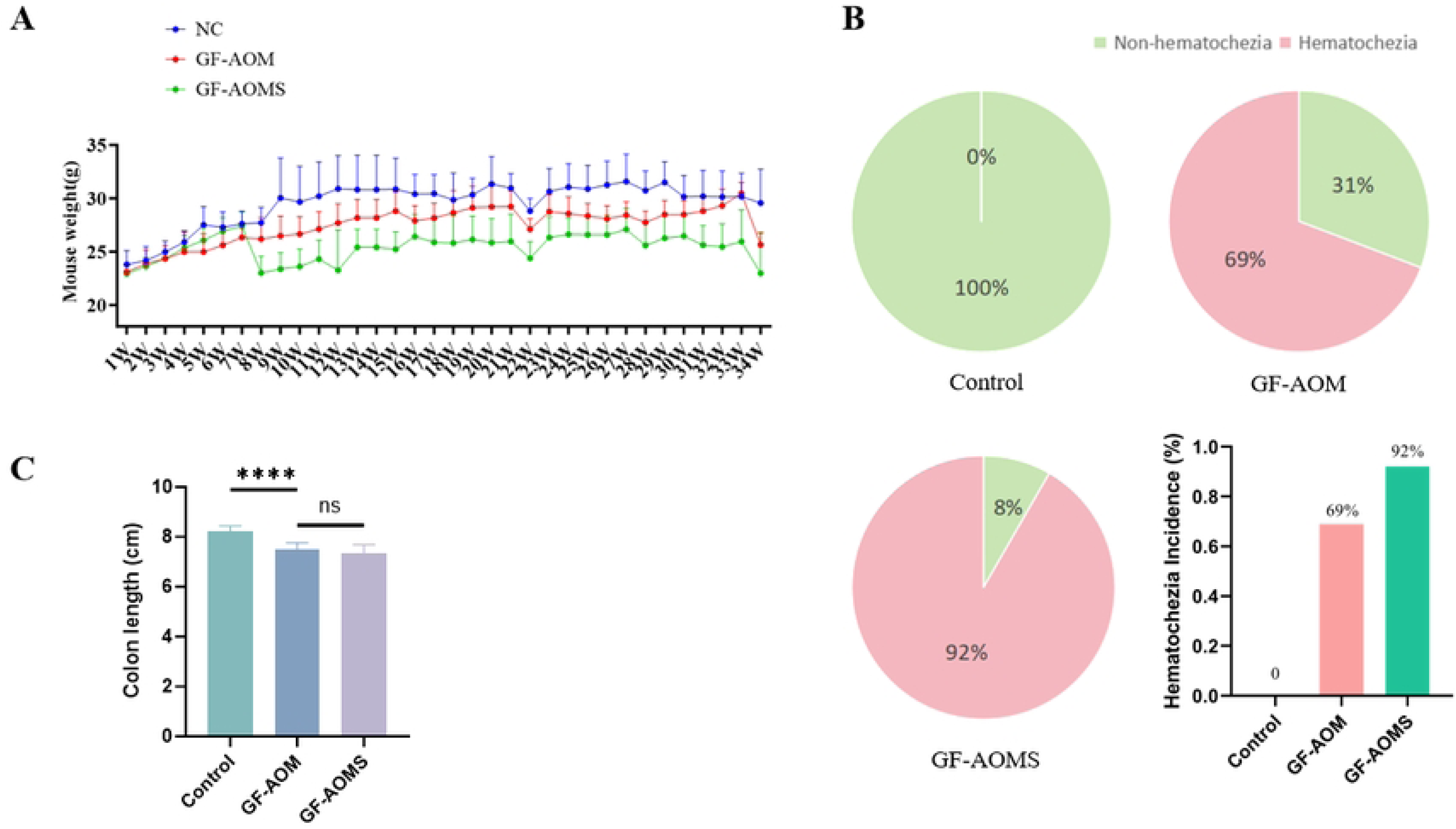
Body weight, fecal blood incidence, and colon length in mice. A: Body weight monitoring of mice. B: Incidence of fecal blood in each group. C: Colon length measurement. Control: control group; GF-AOM: AOM-only intervention group; GF-AOMS: AOM combined with cigarette smoke intervention group.

**Supplementary Figure 2.**
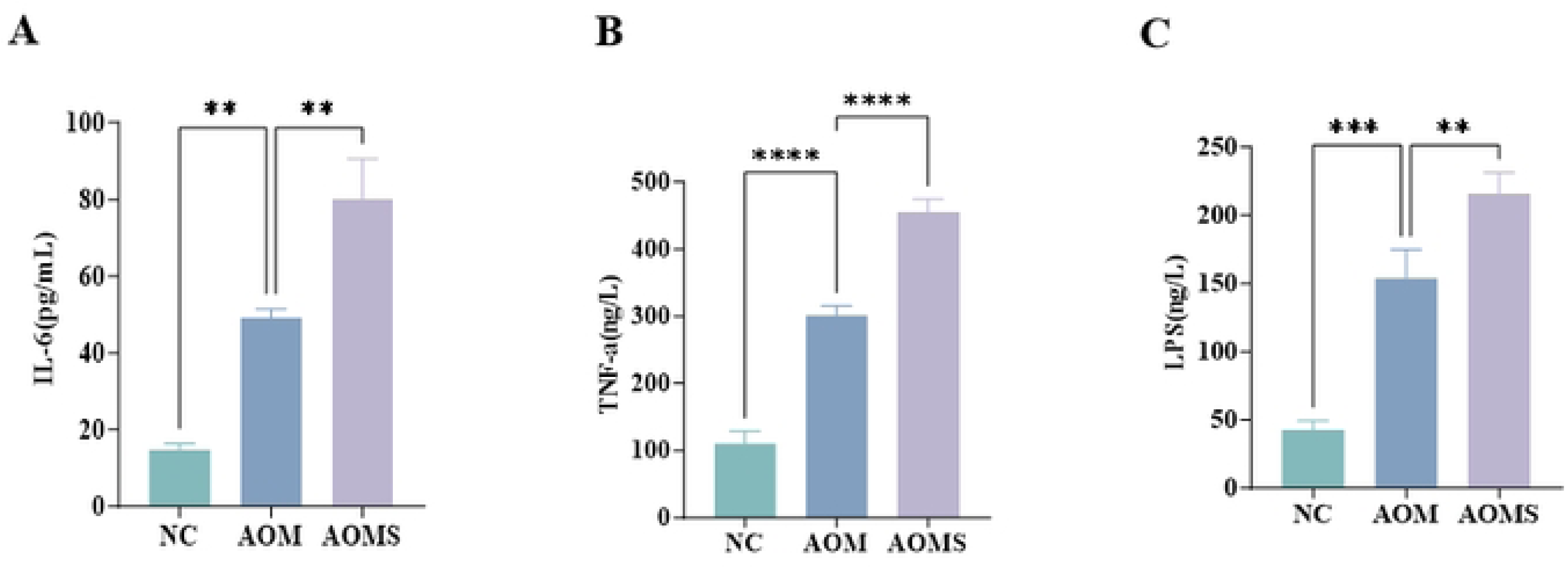
Serum levels of IL-6, TNF-α, and LPS in mice. A: Serum IL-6 levels. B: Serum TNF-α levels. C: Serum LPS levels. NC: control group; AOM: AOM-only intervention group; AOMS: AOM combined with cigarette smoke intervention group. Data are presented as mean ± SEM. Comparisons between two groups were performed using unpaired t-test, and comparisons among multiple groups were analyzed by one-way ANOVA. *ns, p > 0.05; *p < 0.05; **p < 0.01; ***p < 0.001; ****p < 0.0001*.

**Supplementary Figure 3.**
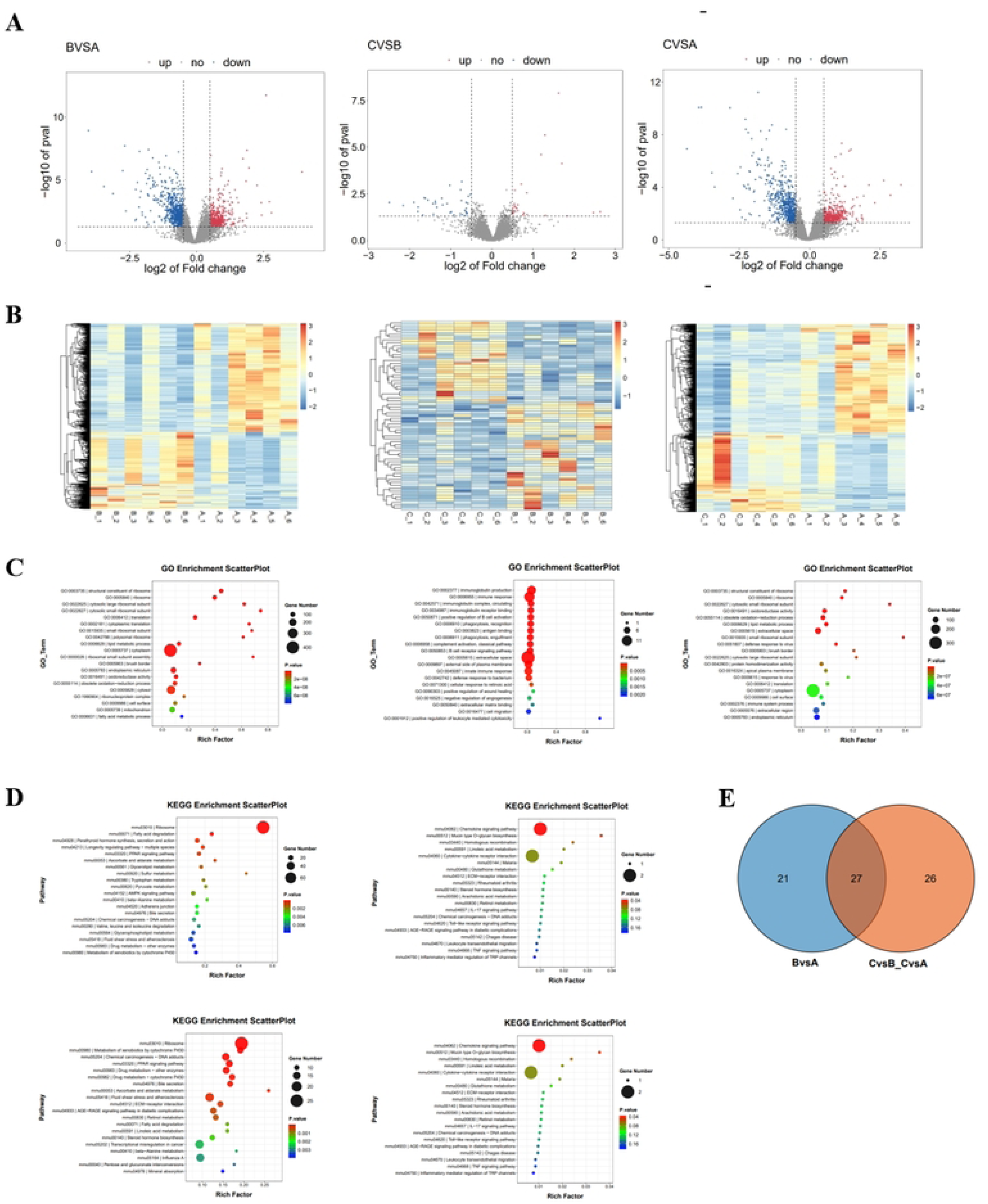
Cigarette smoke regulates 19 signaling pathways in mouse colonic epithelium by affecting 102 genes associated with colorectal cancer. A: Volcano plot of differentially expressed genes (DEGs). Thresholds were set as |log2FoldChange| ≥ 0.5 and p-value < 0.05. Significantly upregulated genes are shown in red, significantly downregulated genes in blue, and non-significant genes in gray. The x-axis represents the fold change of gene expression between groups, and the y-axis represents the statistical significance of expression differences. B: Heatmap of DEGs, where red indicates higher expression and blue indicates lower expression (Top 100 genes with the smallest q-values are shown as an example). C: GO enrichment bubble plot of DEGs. The y-axis represents GO terms, the x-axis represents the enrichment ratio (Rich factor), the bubble color indicates the p-value (darker red indicates higher significance), and the bubble size corresponds to the number of DEGs annotated to each GO term. D: KEGG pathway enrichment scatter plot of DEGs. The x-axis represents the ratio of DEGs annotated to a KEGG pathway over the total DEGs, the y-axis represents KEGG pathways, the bubble size indicates the number of DEGs in each pathway, and the color gradient from blue to red represents increasing significance of enrichment. E: Venn diagram showing shared pathways among comparisons.

**Supplementary Figure 4.**
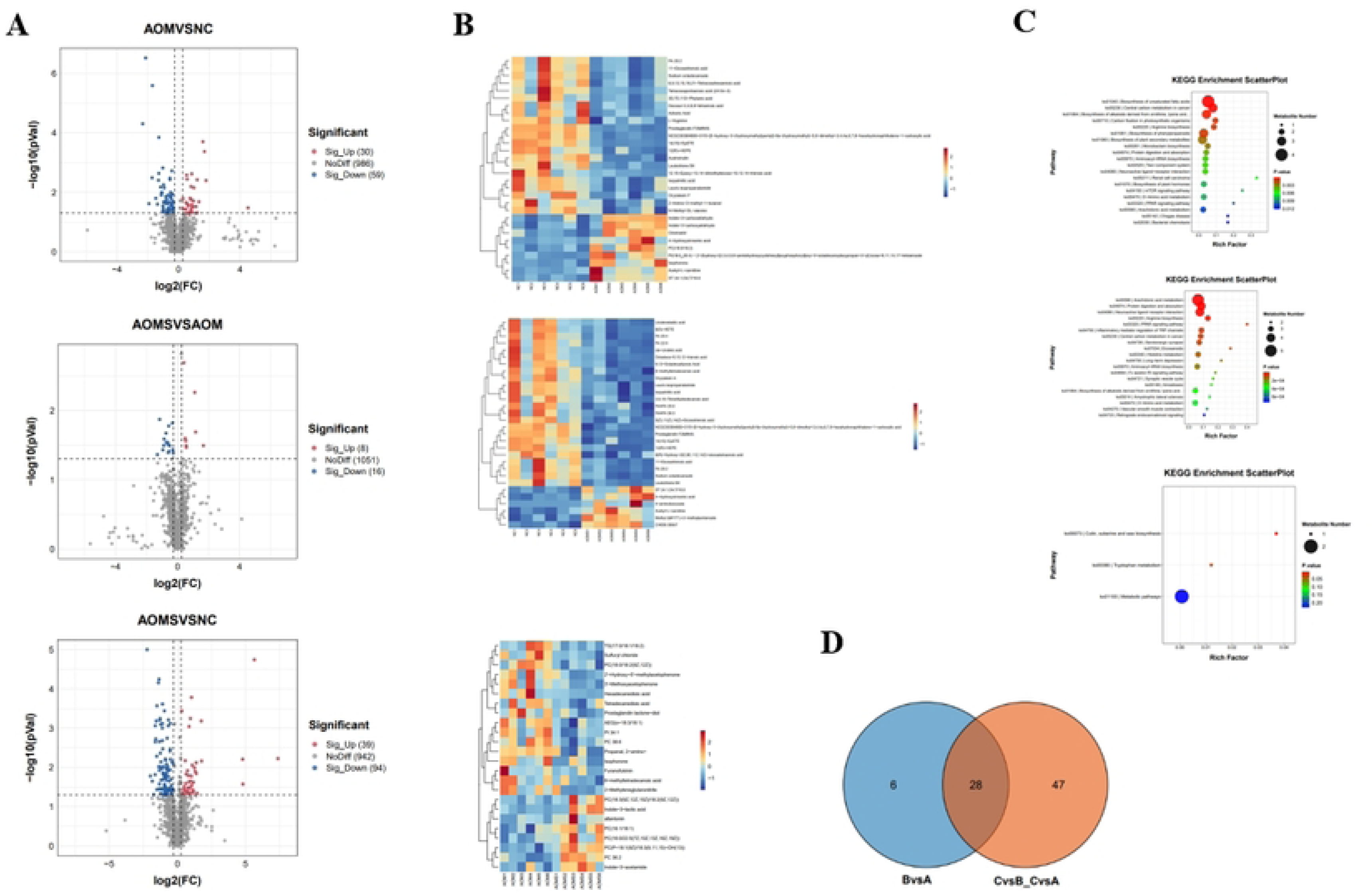
Cigarette smoke alters mouse metabolites associated with colorectal cancer. A: Volcano plot of differential metabolites. The x-axis represents fold change of metabolites between comparison groups, and the y-axis represents -log10(p-value). Significantly upregulated metabolites are shown in red, significantly downregulated metabolites in blue, and non-significant metabolites in gray. B: Heatmap of differential metabolites. Metabolite intensities across samples were normalized and visualized, with color indicating relative abundance. C: KEGG pathway enrichment scatter plot of differential metabolites. The x-axis represents the ratio of differential metabolites annotated to a KEGG pathway over total differential metabolites, the y-axis represents KEGG pathways, bubble size indicates the number of metabolites annotated to each pathway, and color gradient indicates significance. D: Venn diagram showing shared KEGG pathways among comparisons.

**Supplementary Figure 5.**
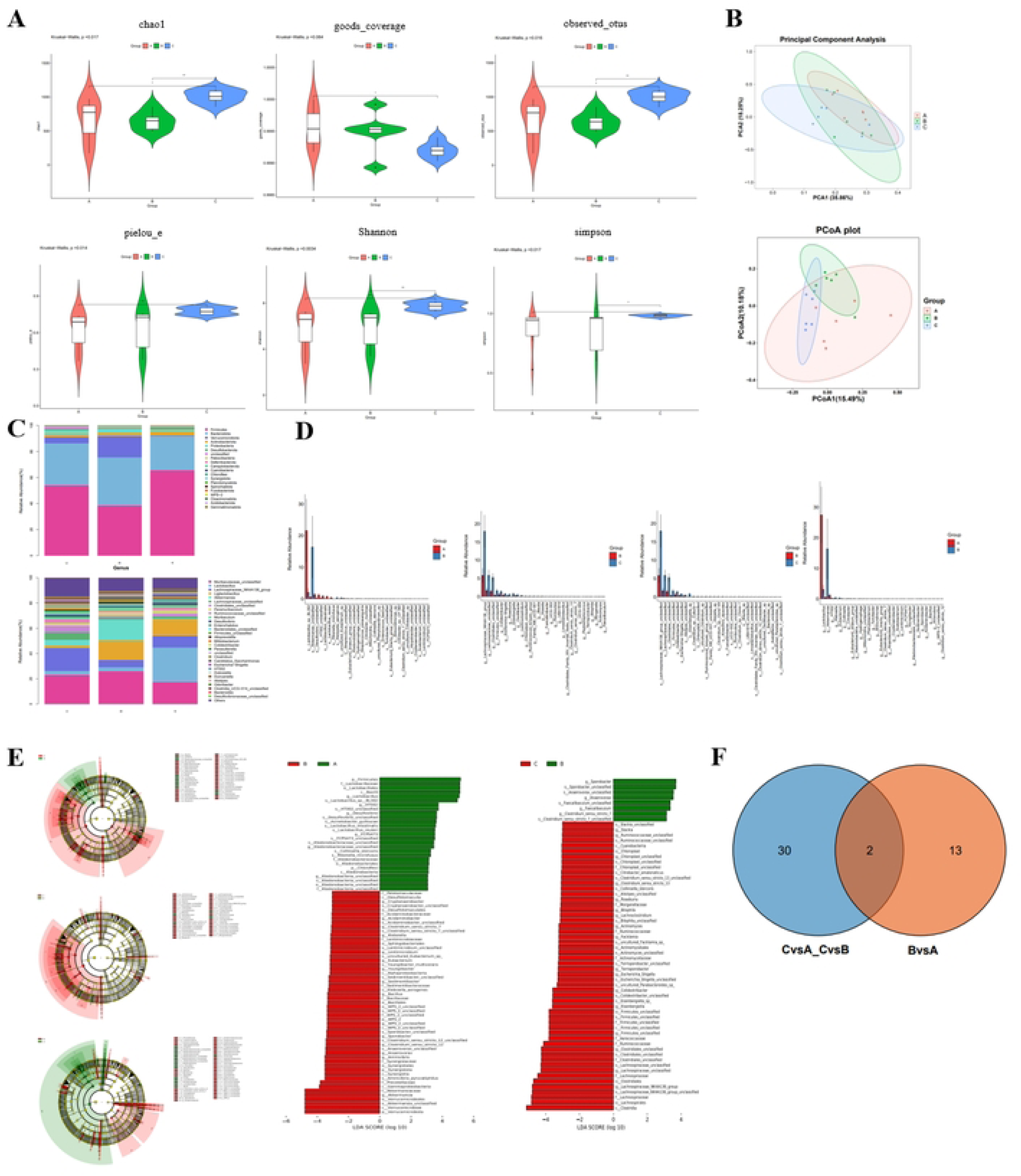
Cigarette smoke alters gut microbiota composition in mice, influencing colorectal cancer development. A: Alpha diversity analysis of gut microbiota. B: Beta diversity analysis of gut microbiota. C: Taxonomic composition at phylum and genus levels. Phylum-level relative abundance is shown on the top, genus-level relative abundance on the bottom. D: Differential abundance analysis of microbial taxa. E: LEfSe analysis of differential taxa. Concentric circles from inner to outer represent seven taxonomic levels: kingdom, phylum, class, order, family, genus, and species. Each node represents a taxon at that level; larger nodes indicate higher abundance. Node color indicates significance: yellow represents no significant difference, red indicates significantly higher abundance in the corresponding group, other colors follow similarly. Significantly differential phyla are directly labeled; other differential taxa are labeled with letters, corresponding to specific species. These results highlight taxa with significant differences across groups and their potential roles under different treatments. F: Venn diagram showing shared microbial taxa among groups.

**Supplementary Figure 6.**
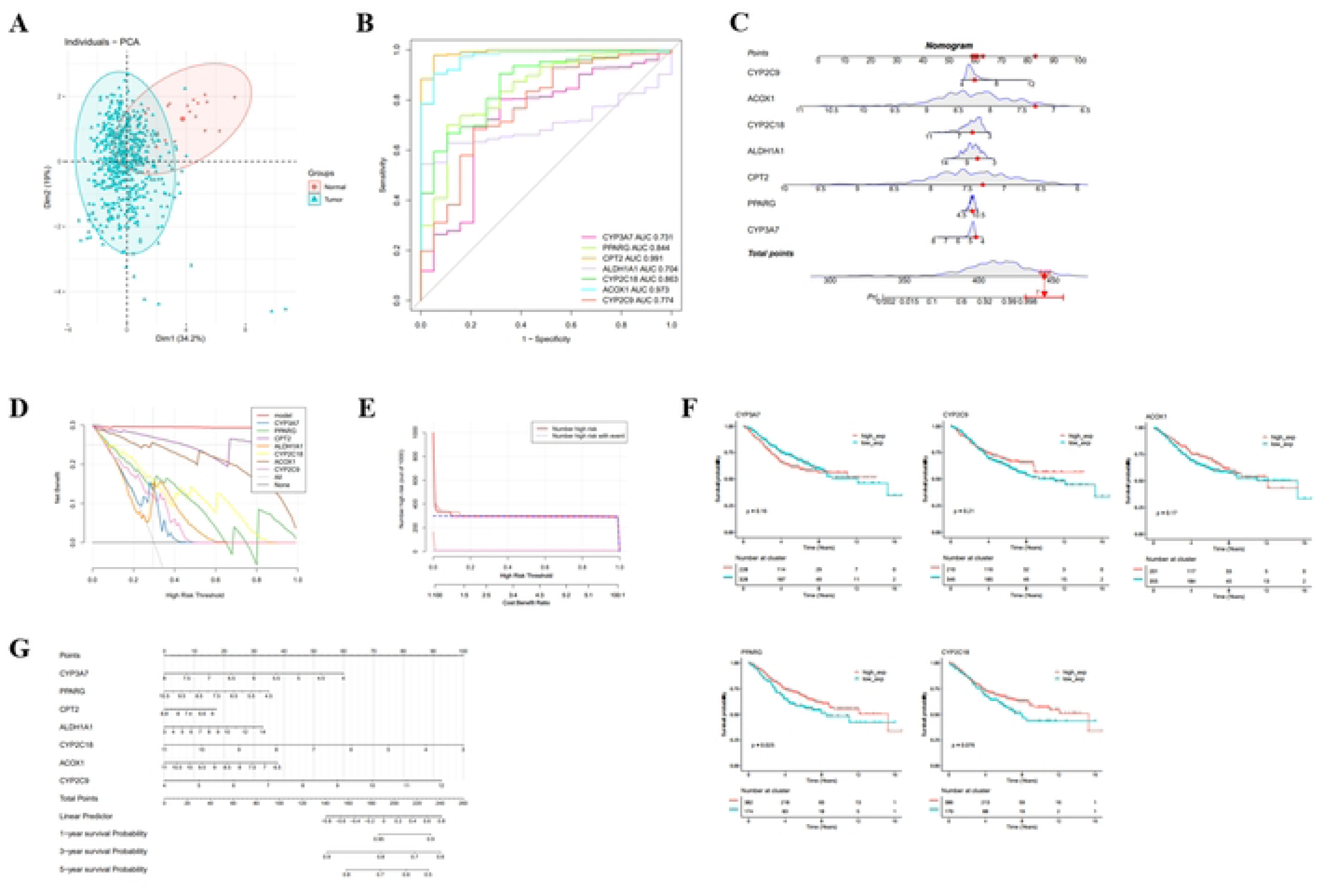
Multi-omics–identified key genes for colorectal cancer diagnostic and prognostic evaluation. A: Principal component analysis (PCA) of candidate genes. B: Receiver operating characteristic (ROC) curves for candidate genes. C: Diagnostic nomogram based on candidate genes. D: Decision curve analysis for diagnostic performance. E: Clinical impact curves of candidate genes. F: Kaplan–Meier survival analysis for CPT2, PPARG, and ALDH1A1. G: Prognostic nomogram for overall survival prediction.

**Supplementary Figure 7.**
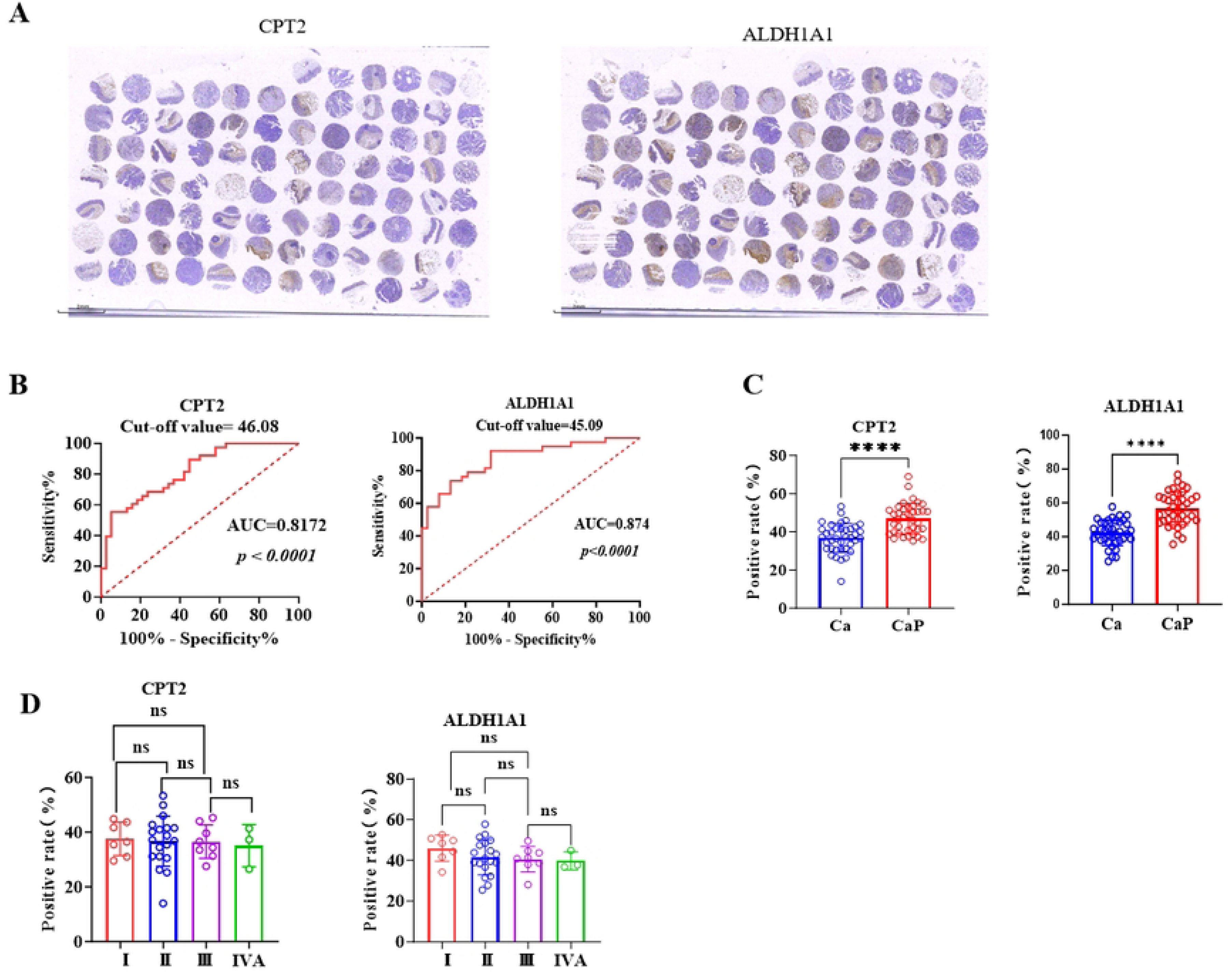
Expression characteristics of CPT2 and ALDH1A1 in colorectal cancer based on cut-off values, AUC, and tissue staging. A: Clinical tissue microarray. B: ROC curves determining the discriminatory ability of CPT2 and ALDH1A1 expression between tumor and adjacent normal tissues. C: Positive expression rates of CPT2 and ALDH1A1 across different TNM stages (I–IVA) of colorectal cancer tissues.

